# Towards understanding diversity, endemicity and global change vulnerability of soil fungi

**DOI:** 10.1101/2022.03.17.484796

**Authors:** Leho Tedersoo, Vladimir Mikryukov, Alexander Zizka, Mohammad Bahram, Niloufar Hagh-Doust, Sten Anslan, Oleh Prylutskyi, Manuel Delgado-Baquerizo, Fernando T. Maestre, Jaan Pärn, Maarja Öpik, Mari Moora, Martin Zobel, Mikk Espenberg, Ülo Mander, Abdul Nasir Khalid, Adriana Corrales, Ahto Agan, Aída-M. Vasco-Palacios, Alessandro Saitta, Andrea C. Rinaldi, Annemieke Verbeken, Bobby P. Sulistyo, Boris Tamgnoue, Brendan Furneaux, Camila Duarte Ritter, Casper Nyamukondiwa, Cathy Sharp, César Marín, Daniyal Gohar, Darta Klavina, Dipon Sharmah, Dong Qin Dai, Eduardo Nouhra, Elisabeth Machteld Biersma, Elisabeth Rähn, Erin K. Cameron, Eske De Crop, Eveli Otsing, Evgeny A. Davydov, Felipe E. Albornoz, Francis Q. Brearley, Franz Buegger, Geoffrey Zahn, Gregory Bonito, Inga Hiiesalu, Isabel C. Barrio, Jacob Heilmann-Clausen, Jelena Ankuda, John Y. Kupagme, Jose G. Maciá-Vicente, Joseph Djeugap Fovo, József Geml, Juha M. Alatalo, Julieta Alvarez-Manjarrez, Kadri Põldmaa, Kadri Runnel, Kalev Adamson, Kari Anne Bråthen, Karin Pritsch, Kassim I. Tchan, Kęstutis Armolaitis, Kevin D. Hyde, Kevin K. Newsham, Kristel Panksep, Adebola A. Lateef, Liis Tiirmann, Linda Hansson, Louis J. Lamit, Malka Saba, Maria Tuomi, Marieka Gryzenhout, Marijn Bauters, Meike Piepenbring, Nalin Wijayawardene, Nourou S. Yorou, Olavi Kurina, Peter E. Mortimer, Peter Meidl, Petr Kohout, R. Henrik Nilsson, Rasmus Puusepp, Rein Drenkhan, Roberto Garibay-Orijel, Roberto Godoy, Saad Alkahtani, Saleh Rahimlou, Sergey V. Dudov, Sergei Põlme, Soumya Ghosh, Sunil Mundra, Talaat Ahmed, Tarquin Netherway, Terry W. Henkel, Tomas Roslin, Vincent Nteziryayo, Vladimir E. Fedosov, Vladimir G. Onipchenko, W. A. Erandi Yasanthika, Young Woon Lim, Nadejda A. Soudzilovskaia, Alexandre Antonelli, Urmas Kõljalg, Kessy Abarenkov

## Abstract

Fungi play pivotal roles in ecosystem functioning, but little is known about their global patterns of diversity, endemicity, vulnerability to global change drivers and conservation priority areas. We applied the high-resolution PacBio sequencing technique to identify fungi based on a long DNA marker that revealed a high proportion of hitherto unknown fungal taxa. We used a Global Soil Mycobiome consortium dataset to test relative performance of various sequencing depth standardization methods (calculation of residuals, exclusion of singletons, traditional and SRS rarefaction, use of Shannon index of diversity) to find optimal protocols for statistical analyses. Altogether, we used six global surveys to infer these patterns for soil-inhabiting fungi and their functional groups. We found that residuals of log-transformed richness (including singletons) against log-transformed sequencing depth yields significantly better model estimates compared with most other standardization methods. With respect to global patterns, fungal functional groups differed in the patterns of diversity, endemicity and vulnerability to main global change predictors. Unlike α-diversity, endemicity and global-change vulnerability of fungi and most functional groups were greatest in the tropics. Fungi are vulnerable mostly to drought, heat, and land cover change. Fungal conservation areas of highest priority include wetlands and moist tropical ecosystems.

## Introduction

Human activities affect nearly all habitats through changes in climate and land-use, which in turn alter vegetation cover and composition. These changes negatively impact many species that have narrow environmental tolerances and limited dispersal capacity across anthropogenic landscapes (Schulte to Bühne et al. 2020). Anthropogenic impacts most strongly affect endemic species – i.e., taxa with small distribution ranges and narrow ecological niches (Brook et al. 2008). Diversity of endemic plants and animals is higher in areas characterized by historical stability, high precipitation, environmental heterogeneity, and insularity. Unfortunately, these areas usually coincide with major human degradations of the environment (Kier et al. 2009; Stein et al. 2014; Sandel et al. 2020).

Unlike the situation with plants and animals, global patterns of fungal diversity, endemism and vulnerability to environmental change remain virtually unknown (Cameron et al. 2019; Guerra et al. 2021b; but see Talbot et al. 2014; Davison et al. 2015). This is alarming, given the fundamental roles that fungi play in carbon and nutrient cycling processes (Wardle & Lindahl 2014; Crowther et al. 2019). Comparative studies have indicated that aboveground and belowground biodiversity are driven by different environmental predictors at local and global scales (Cameron et al. 2019; Le Provost et al. 2021). This suggests differential responses of macro- and microorganisms to land use and climate changes (Guerra et al. 2021b). As for plants and animals, soil fungal communities are likely vulnerable to global change drivers. For instance, high-temperature stress (Malcolm et al. 2008; Barcenas-Moreno et al. 2009; Morgado et al. 2015; Misiak et al. 2021) and prolonged drought (Schmidt et al. 2017; de Vries et al. 2018) can alter fungal growth, functionality and community composition. Likewise, changes in land use that result in habitat fragmentation may lead to shifts in prevalence of pathogenic, mutualistic, and free-living fungal groups (Brinkmann et al. 2019; Makiola et al. 2019; Le Provost et al. 2021; Rodriguez-Ramos et al. 2021).

While thousands of plant and animal species are listed as threatened on the IUCN global Red List, only 262 out of an estimated 2.2-3.8 million fungal species (Hawksworth & Lücking 2017) have been listed as such. The majority of these are from high-income countries in temperate regions (IUCN 2021) and are from fungal groups that make conspicuous macroscopic fruiting bodies (Cui et al. 2021). However, the vast majority of fungi produce no or inconspicuous fruiting bodies and are therefore hard to survey, which has hampered their conservation assessment (Gonçalves et al. 2021).

Here we used the most advanced high-resolution sequencing technology to globally survey soil fungal diversity and assess their endemicity and vulnerability to global change. We hypothesized that i) the endemicity of fungi is relatively higher in the tropics due to greater regional climatic stability; and ii) vulnerability of fungi to global change is highest in habitats experiencing the strongest global warming effects (polar regions) and intensive land use (dry tropics). We predicted that because of their intimate associations with other organisms, endemicity and vulnerability patterns are more evident for macrofungi and biotrophic groups compared with saprotrophic microfungal groups. We then propose global conservation priorities for these ecologically pivotal fungi.

## Results and Discussion

### Fungal diversity

We used the recently generated Global Soil Mycobiome consortium dataset (GSMc; 3,200 plots, Tedersoo et al. 2021b) along with data from five other global soil surveys (**Fig. 1**; see methods) and international nucleotide sequence databases to determine the diversity and endemicity of fungal functional groups – *viz*. arbuscular mycorrhizal (AM) fungi, ectomycorrhizal (EcM) fungi, non-EcM Agaricomycetes (mostly saprotrophic macrofungi), molds, pathogens, opportunistic human parasites (OHPs, mostly thermophilic saprotrophs), early-diverging unicellular lineages (mostly chytrids, aphelids, and rozellids), and yeasts. Compared to previous meta-analytical approaches (e.g. Vetrovsky et al. 2019), our cumulative data comprise the largest available globally standardized database based on directly comparable soil sampling and long-read molecular analysis protocols. Collectively, all datasets yielded 20,182,427 fungal reads composed of 905,841 ‘species’ – operational taxonomic units (OTUs), each defined as <98% sequence similarity of the rRNA ITS barcode from all other OTUs. The genera *Tomentella* (Basidiomycota), *Penicillium* (Ascomycota), and *Mortierella* (Mortierellomycota) were the most species-rich (**Fig. 1**).

**Fig. 1.**
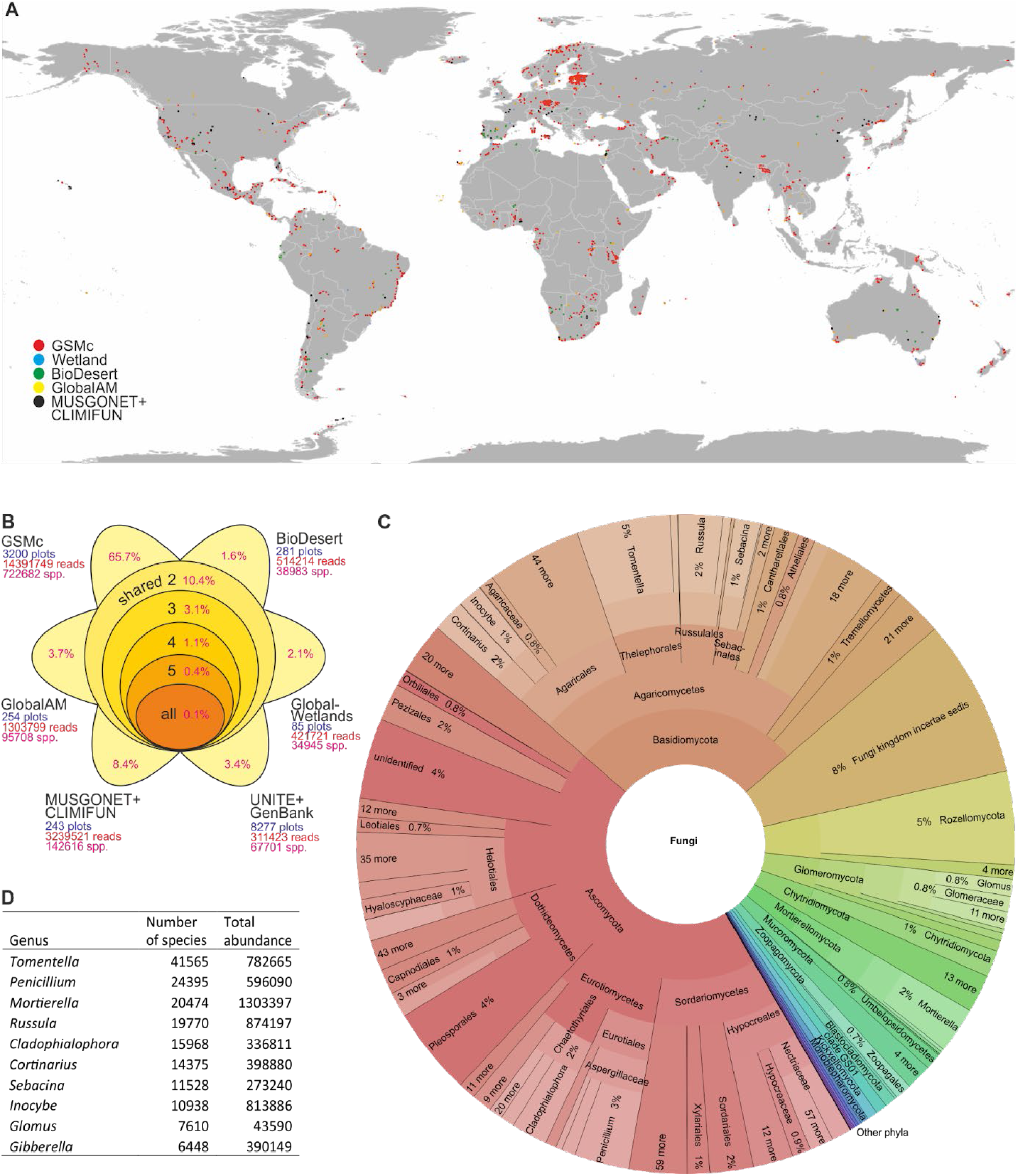
Distribution of samples and fungal species across datasets. (A) Global sampling map, with different symbols representing different datasets; (B) species distribution of fungi among datasets, with the proportion of unique and shared species indicated in the diagram; (C) Krona chart indicating taxonomic distribution of fungal species (interactive chart can be browsed at https://plutof.ut.ee/#/doi/10.15156/BIO/2483900); (D) species richness and total read abundance of the top 10 most diverse fungal genera.

We combined machine-learning and general linear modeling (GLM) approaches to find the best predictors of fungal species richness and the Shannon index of diversity for settling the contrasting results obtained from previous global studies (Tedersoo et al. 2014; Egidi et al. 2019; Vetrovsky et al. 2019). At the site scale (α-diversity), the best supported results were obtained for residuals of logarithm-transformed richness accounting for sequencing depth (**Fig. 2**). Since the datasets were retrieved using different sampling design and therefore differed strongly in the inferred richness (**Fig. 3**), we focused mainly on analyses of the largest, GSMc dataset. Total fungal richness had a broadly unimodal relationship with soil pH (R^2^_adj_=0.133) and responded positively to vegetation age (R^2^_adj_ = 0.045; **Fig. 4**). Deserts and Antarctic habitats supported the lowest richness among all biomes (**Fig. 4**). We validated the results using several datasets, in which fungal richness had a unimodal relationship with soil pH and positive response to mean annual precipitation (MAP)(**Fig. 5**). Across datasets, fungal γ-diversity at the ecoregion level was best explained by average MAP (R^2^_adj_=0.179; **Fig. 6**). The differences in richness trends between α-diversity and γ-diversity indicate that high precipitation favors niche differentiation at the regional scale, as reflected by higher turnover between sites (i.e. increasing α-to γ-diversity).

**Fig. 2.**
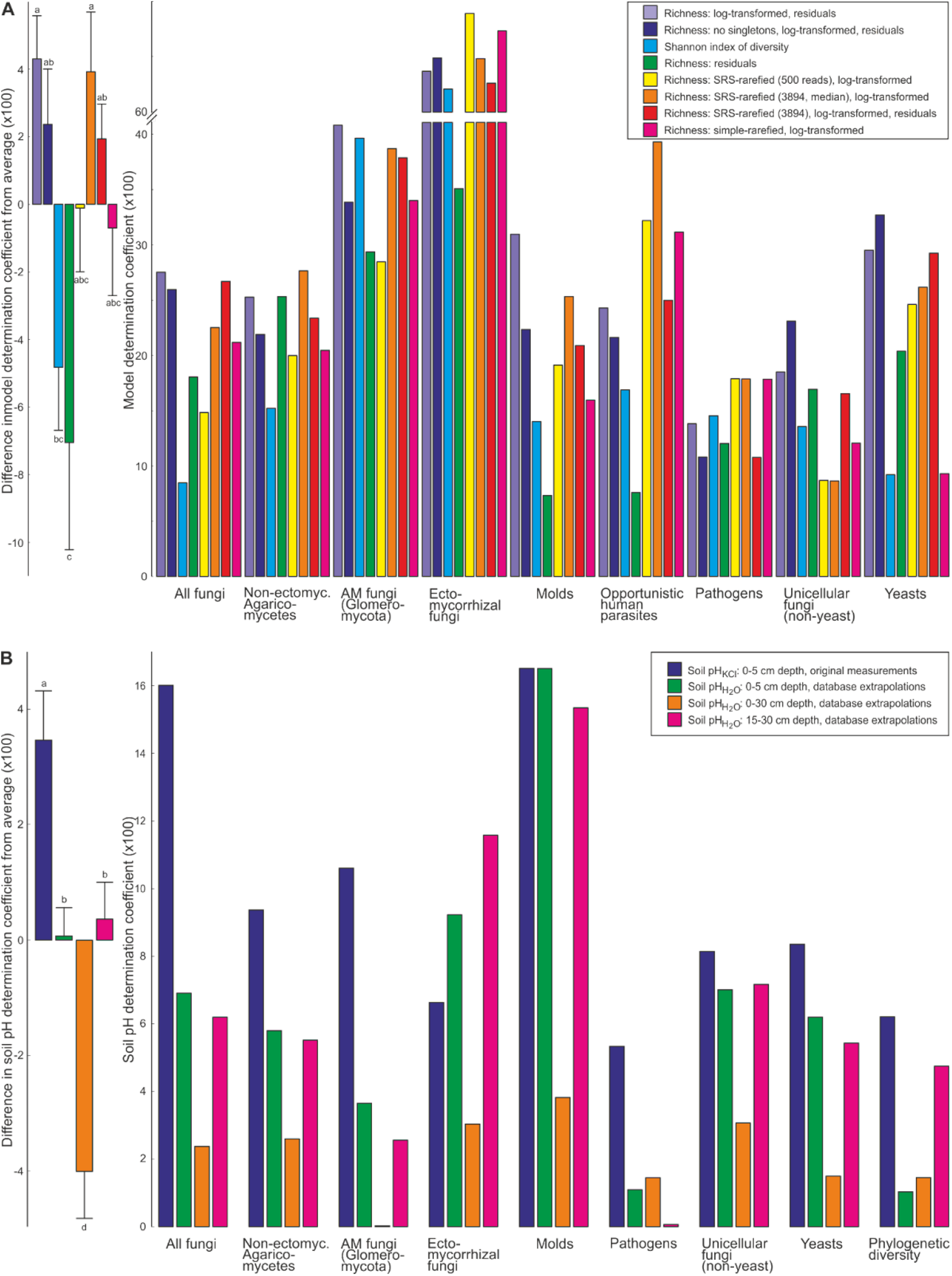
Comparison of (A) richness proxies (use of log-transformation, residuals of sequencing depth, SRS or simple rarefaction) and (B) measures of soil pH on analytical performance. Relative goodness was estimated based on the determination coefficients of the best models (A) or pH-only models (B). In left panels, significant among-group differences are indicated with different letters based on Tukey Posthoc tests; bars, means; whiskers, SE. Soil pH_KCl_ were determined experimentally, whereas pH_H2O_ were obtained from Poggio et al. (2021).

**Fig. 3.**
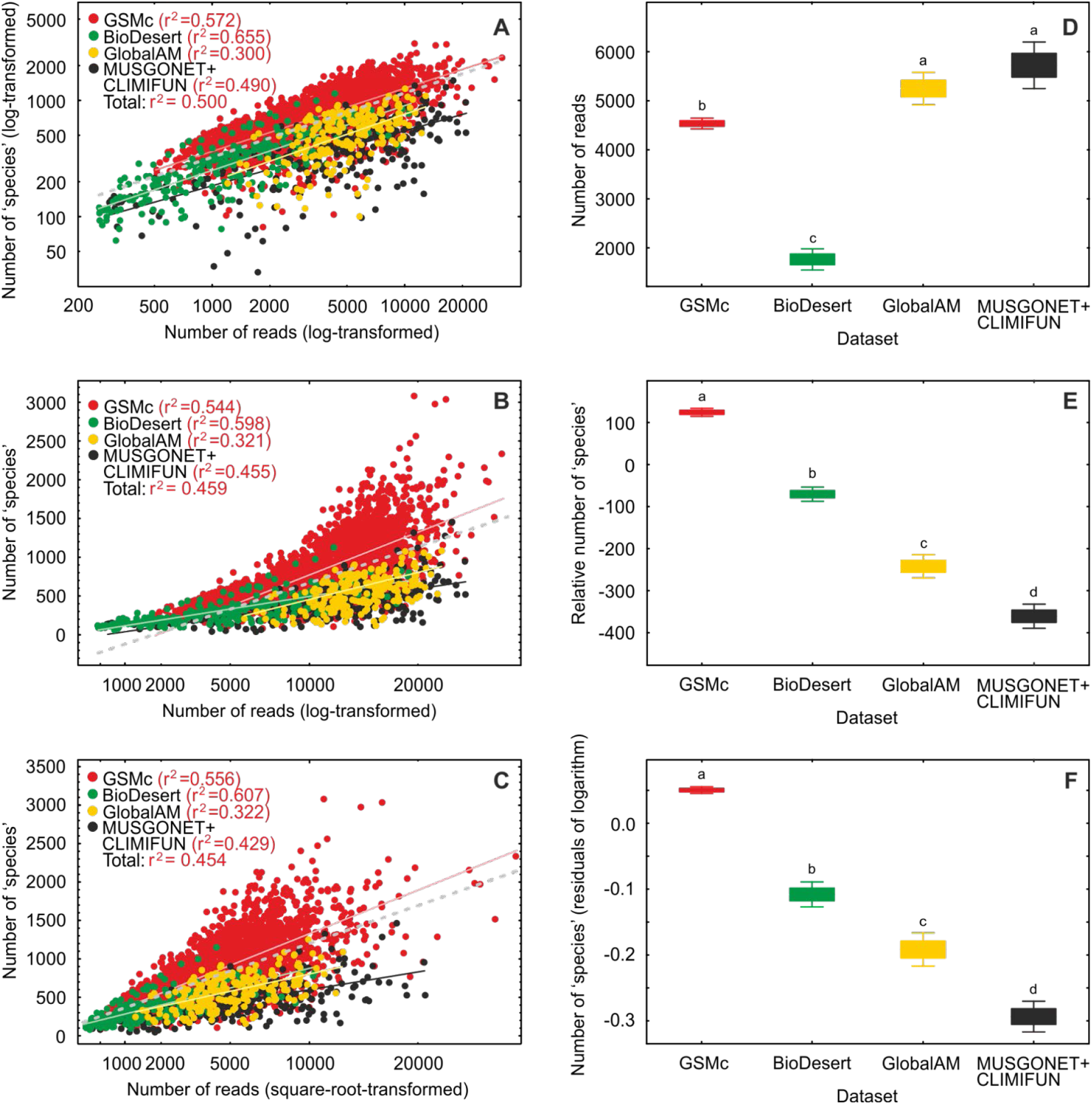
Relative ‘species’ accumulation curves (A-C), sequencing depth (D) and ‘species’ richness (E-F) across four datasets. (A) The log-log relationship between the number of reads and ‘species’ richness that was used for calculation of residuals and further analyses; (B-C) Relatively lower performance of log-linear relationships of log-transformed and square-root-transformed sequencing depth; (D) Initial differences in sequencing depth among datasets; (E-F) Fungal ‘species’ richness differences relative to the average in the raw data (F) and residuals of the log-log regression analysis (F). In D-F, boxes indicate standard errors around the mean and whiskers indicate 95% confidence intervals; letters above whiskers indicate statistically significant differences among datasets (using log-transformed data for D-E). These analyses indicate that the log-log transformation for calculating residuals is relatively more robust compared with other methods and that richness estimates from studies with different methods cannot be directly compared.

**Fig. 4.**
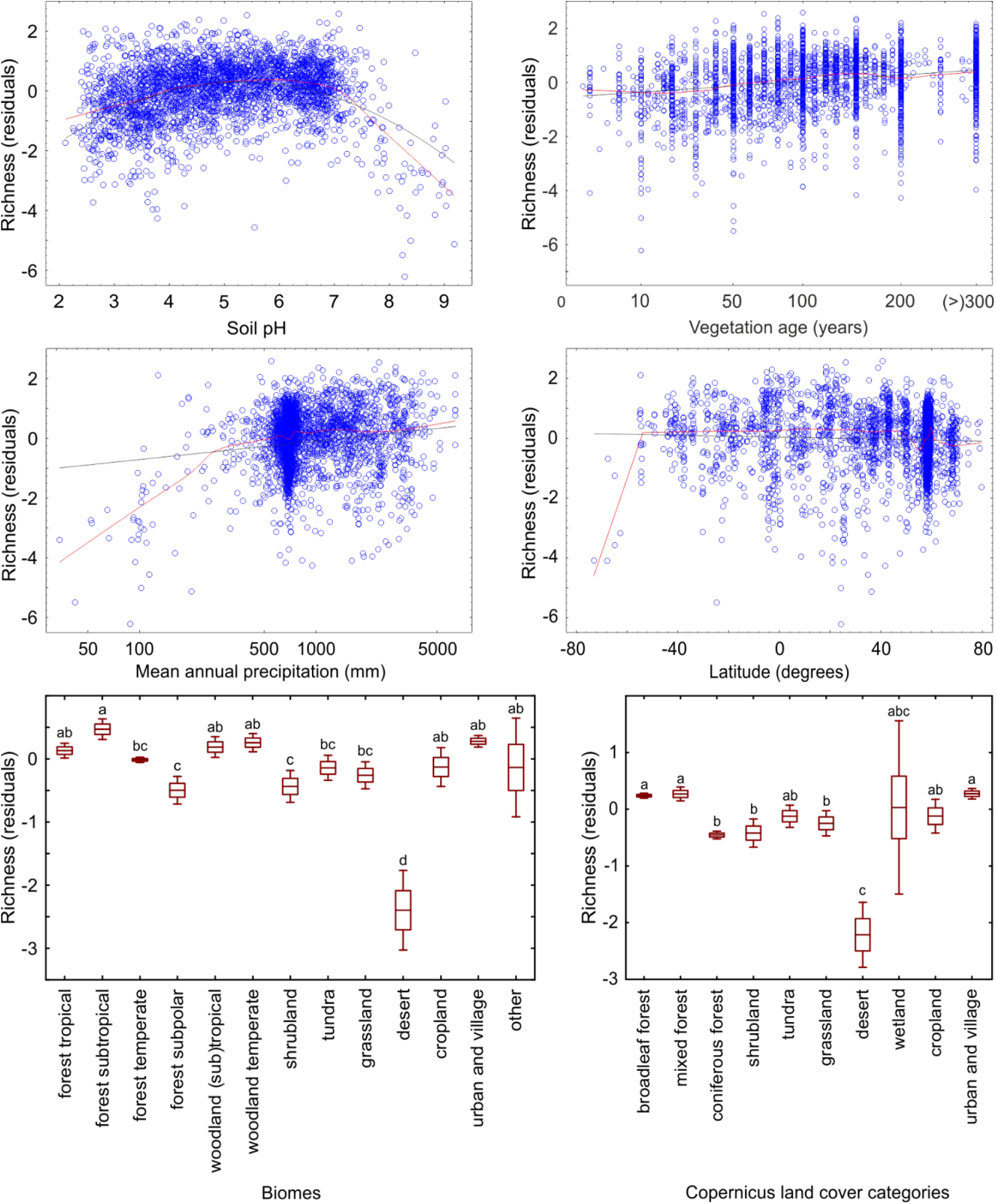
Response of α-diversity of all fungi to soil pH, vegetation age, mean annual precipitation, latitude, biomes, and land cover categories. For continuous predictors, black lines indicate linear and polynomial fits and red lines indicate lowess fits. For categorical predictors, boxes represent standard error around the mean (central line), whiskers depict 95% CI and letters above boxes indicate statistically significant different groups (P<0.001).

**Fig. 5.**
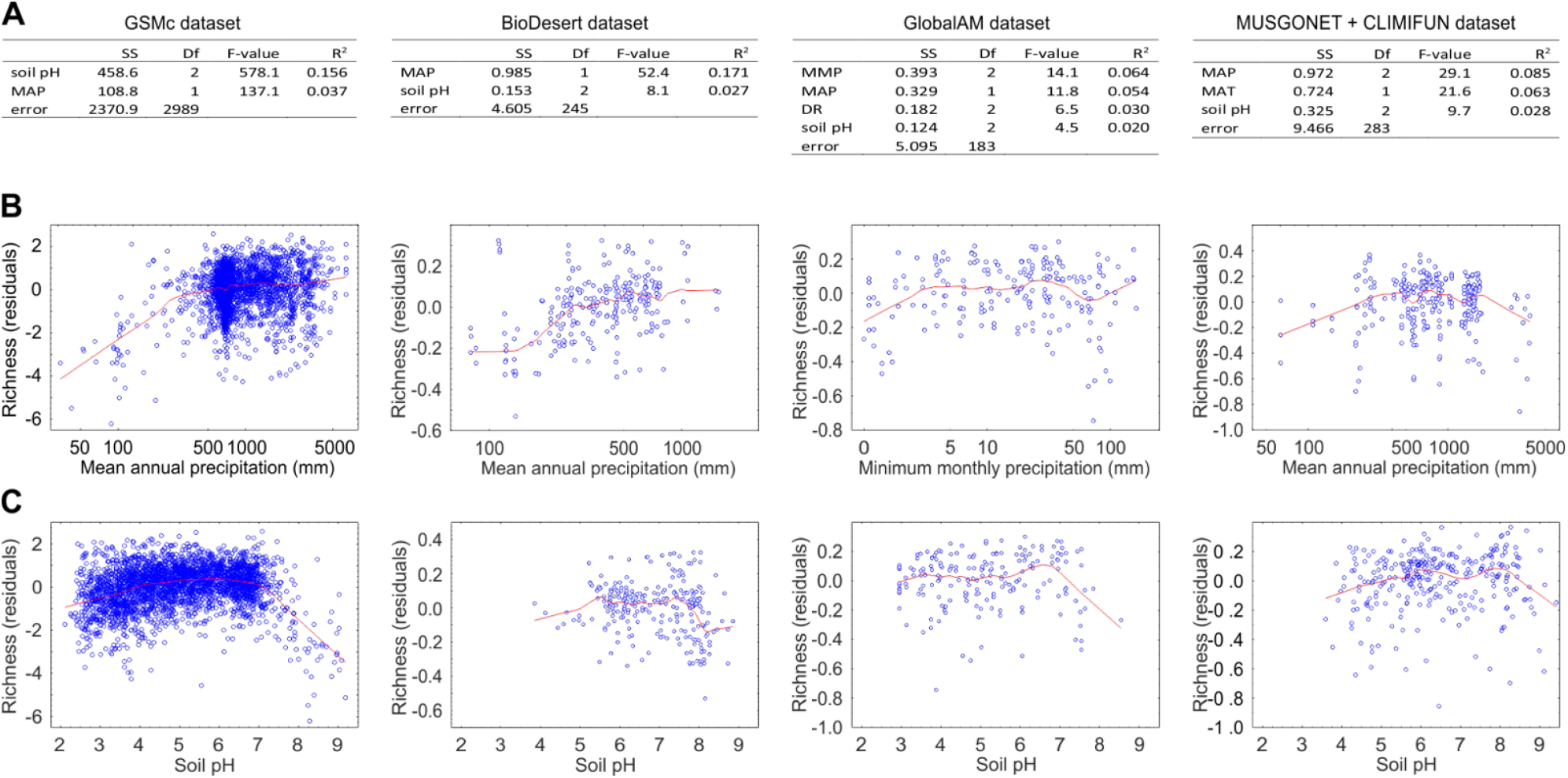
Comparison of α-diversity patterns in all fungi across four most inclusive datasets: (A) best models (only bioclimatic variables and soil pH were included in model selection), (B) lowess regression curves for the best-fitting climatic predictor, and (C) lowess regression curves for soil pH.

**Fig. 6.**
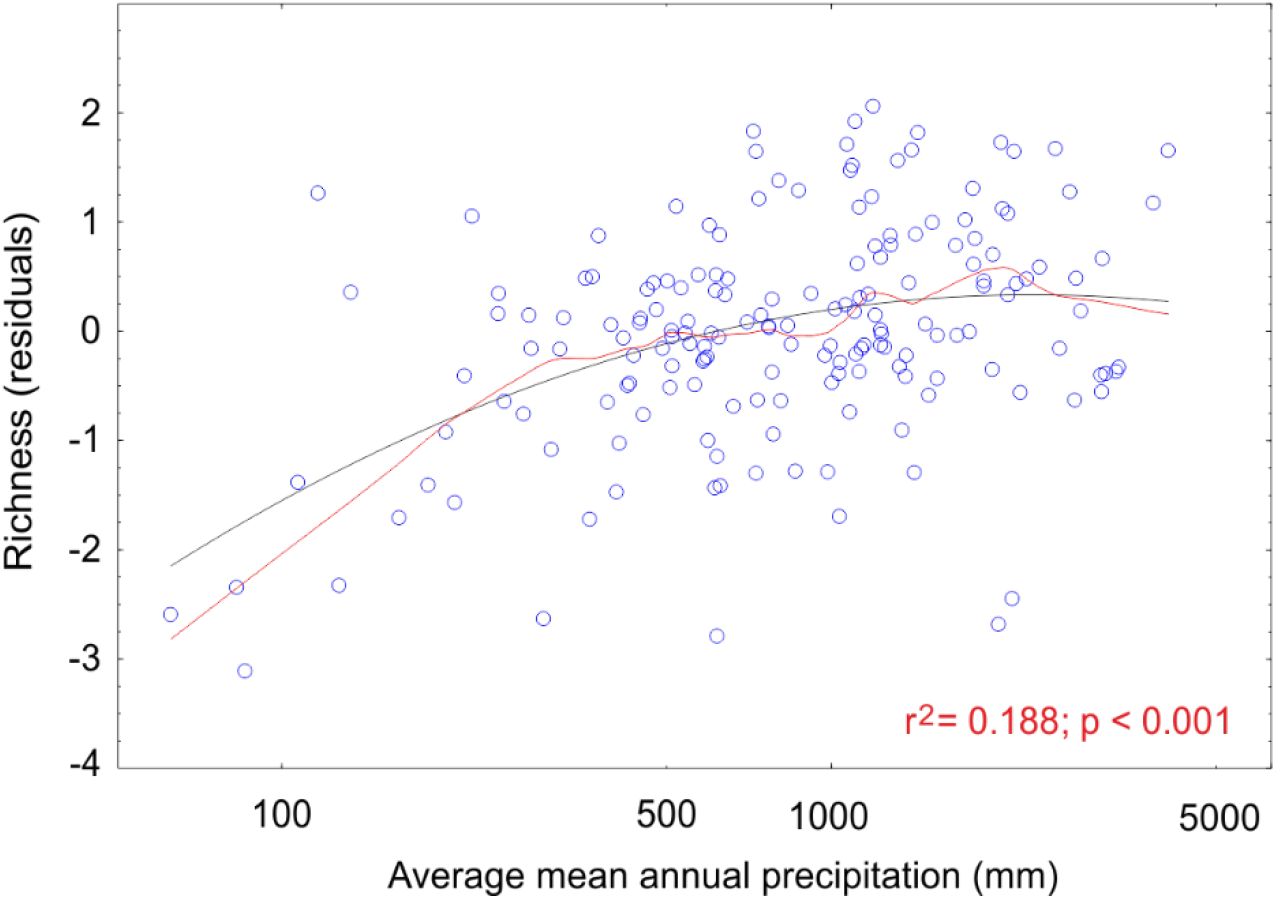
The effect of average mean annual precipitation on γ-diversity of fungi at the ecoregion scale. Black line, best quadratic fit; red line, lowess curve.

Our results thus update previous patterns of α-diversity decrease (Tedersoo et al. 2014) or increase (Vetrovsky et al. 2019) at high latitudes and confirm relatively lower fungal diversity in Antarctica. The latter pattern has been/can be ascribed to low plant diversity and coverage (Newsham et al. 2016). The more prominent latitudinal gradient in γ-diversity reflects a greater positive effect of MAP on the regional fungal species pool. Disregarding Antarctica, the lack of a global α-diversity latitudinal gradient in fungi is unique among terrestrial organisms (Kinlock et al. 2018). By comparison, the γ-diversity patterns detected resemble those found for soil fauna (Aslani et al. 2022; Potapov et al. 2022) and protistan parasites (Oliverio et al. 2020), all which show slight richness peaks in tropical latitudes. The distinctly weaker latitudinal diversity gradients of soil organisms compared with most aquatic and terrestrial macro-organisms may be related to indirect effects of temperature-related climatic variables as well as soil pH and C/N ratio as main drivers of soil habitat quality. The differences may also be related to higher dispersal capacity of soil organisms who have microscopic body sizes or dispersal propagules (Soininen et al. 2013; Aslani et al. 2022).

### Fungal endemicity

To estimate relative endemism among the world’s ecoregions **(Fig. 7; Table 1; see methods**), we combined indices of community similarity, uniqueness, and species ranges into an overall endemicity index (see Methods). Five metrics were combined, including the number and proportion of endemic species, mean maximum geographical range of species, Jaccard index, and beta-sim index (**Box 1**). We found that endemicity of all fungi peaked in moist tropical biomes and it was positively related to mean annual air temperature (MAT; R^2^_adj_=0.277; **Fig. 8**) and soil acidity (R^2^_adj_=0.108; **Table 2**).

**Table 1.**
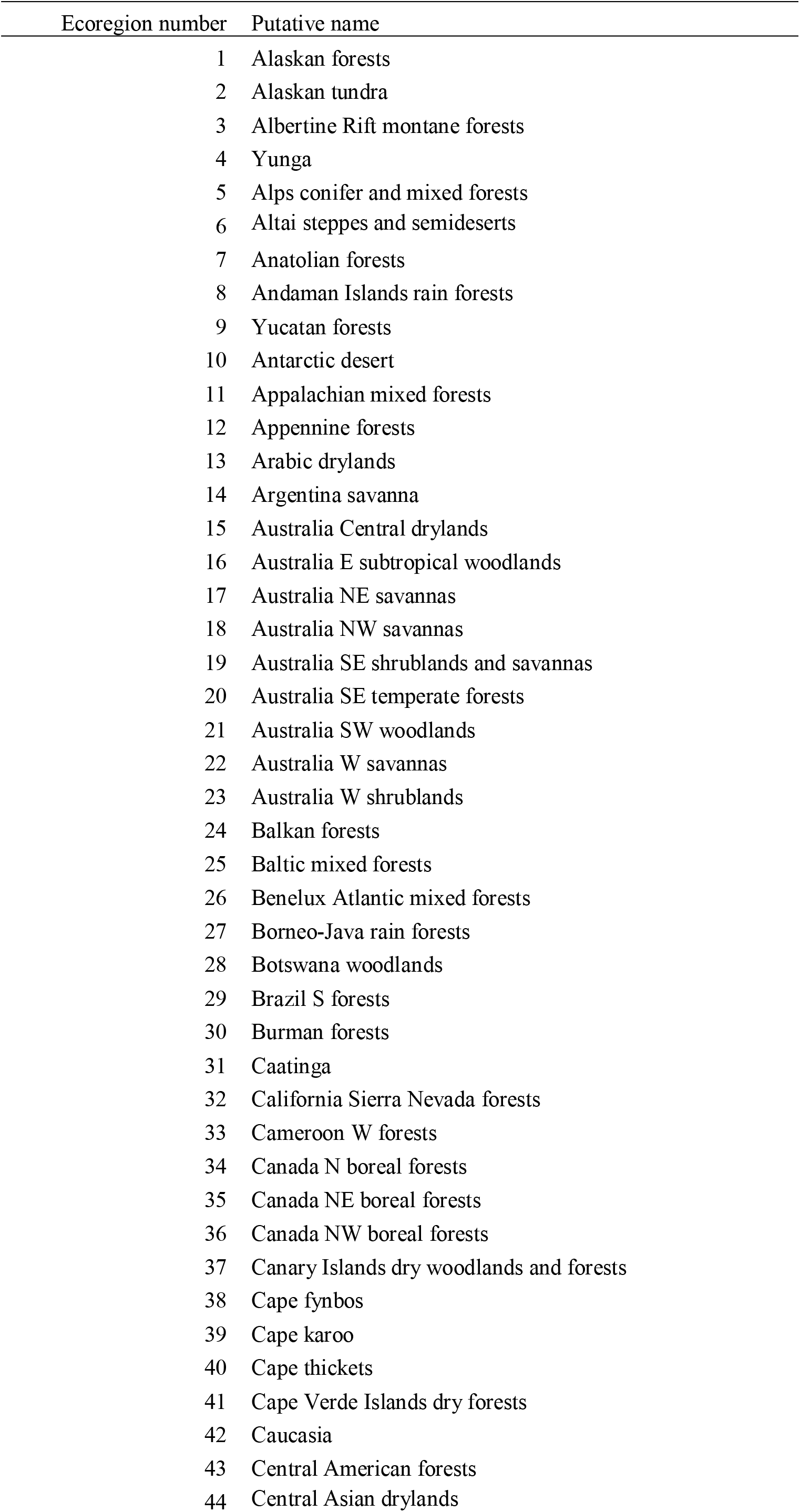
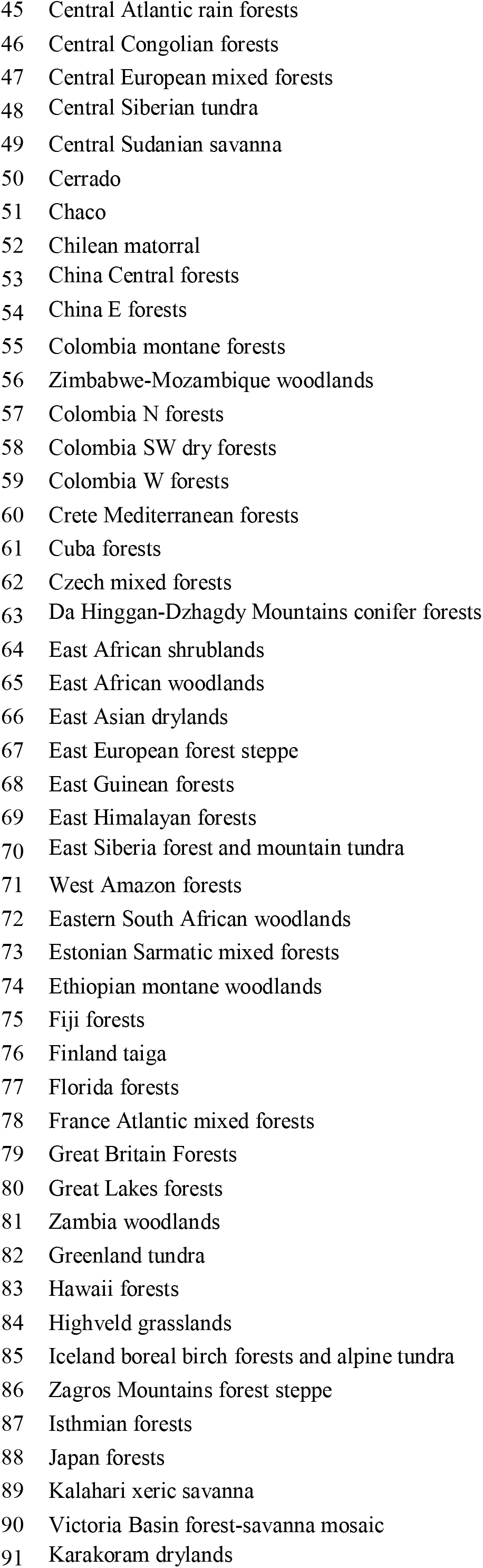
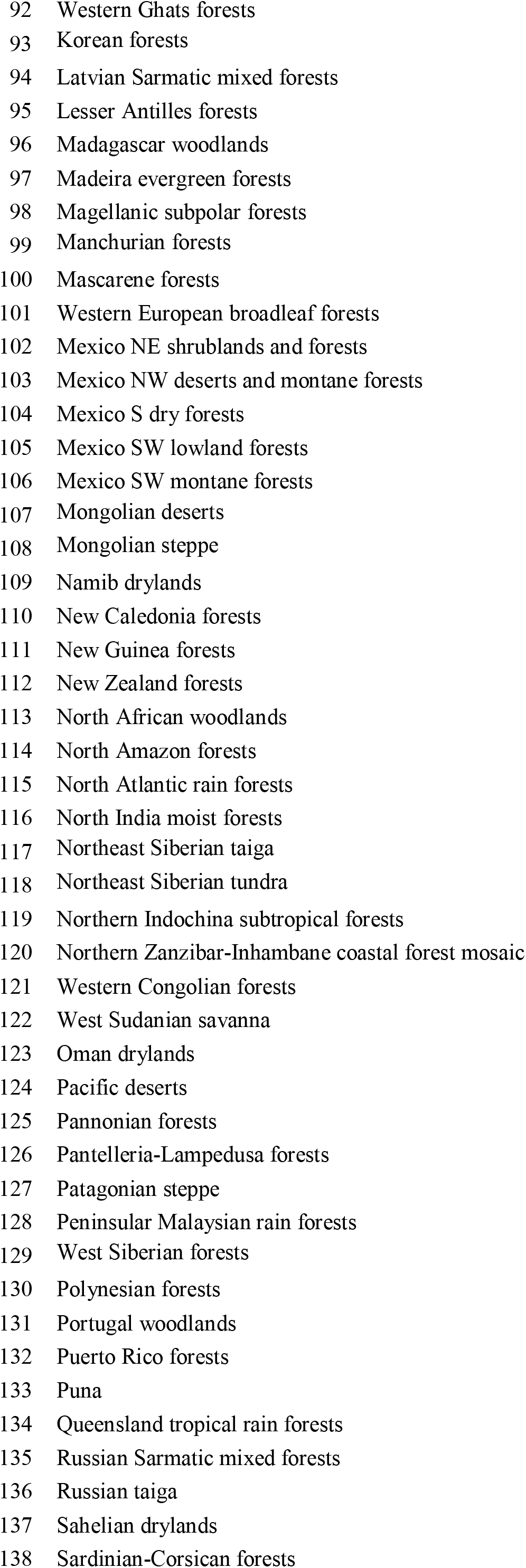
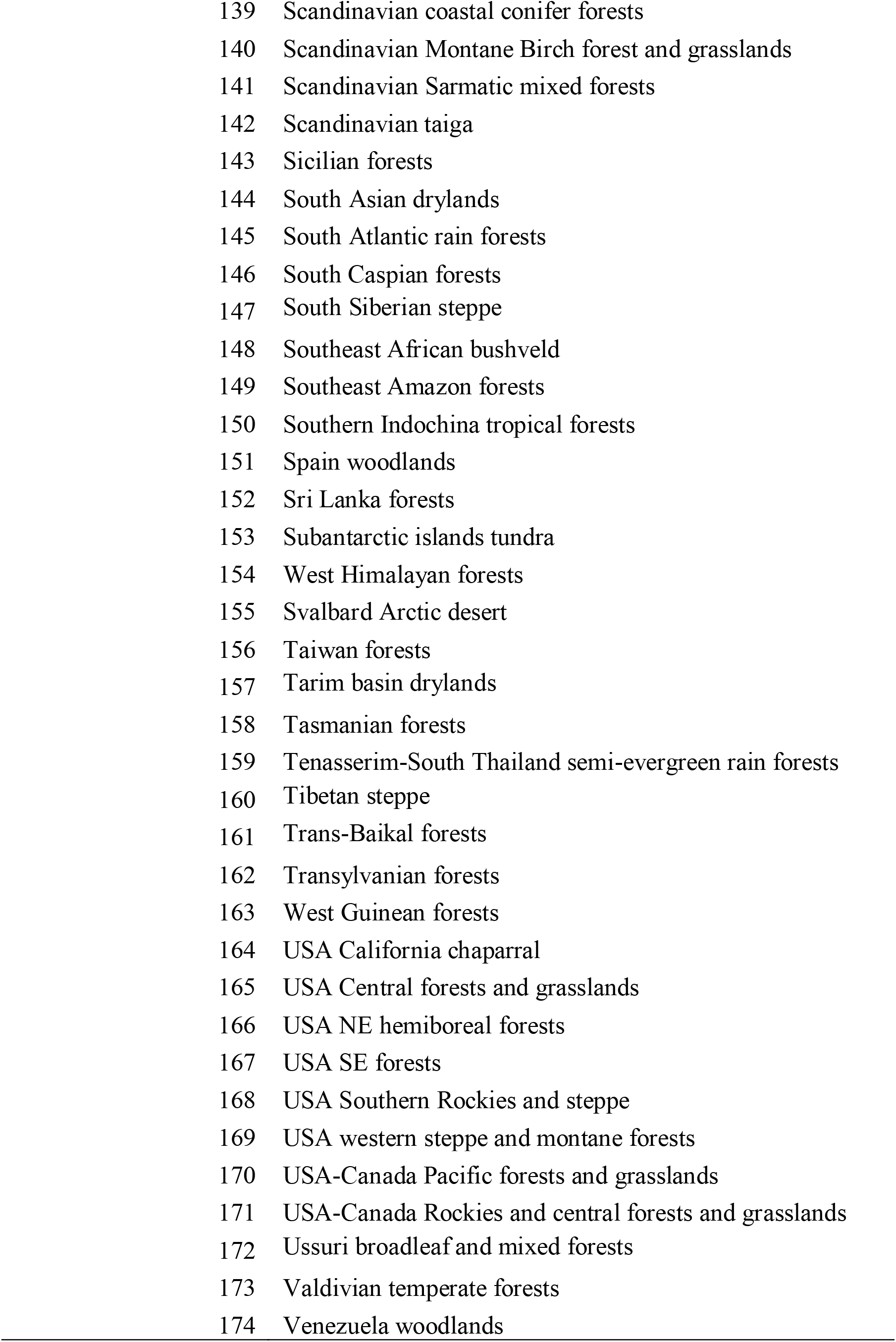
The ecoregions used in endemicity analyses.

**Table 2.**
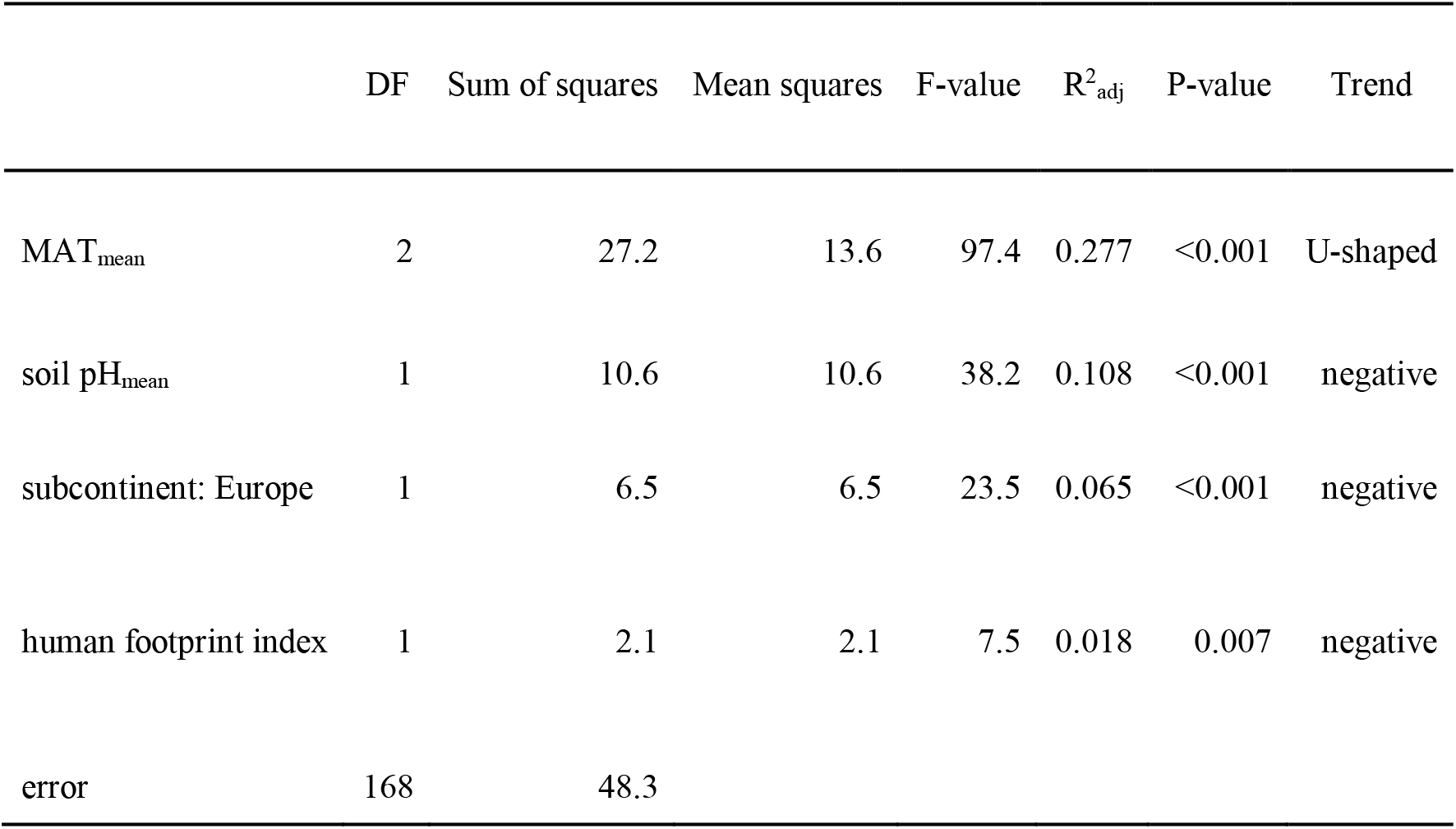
The best predictors of endemicity indices in ecoregions for all fungi.

**Fig. 7.**
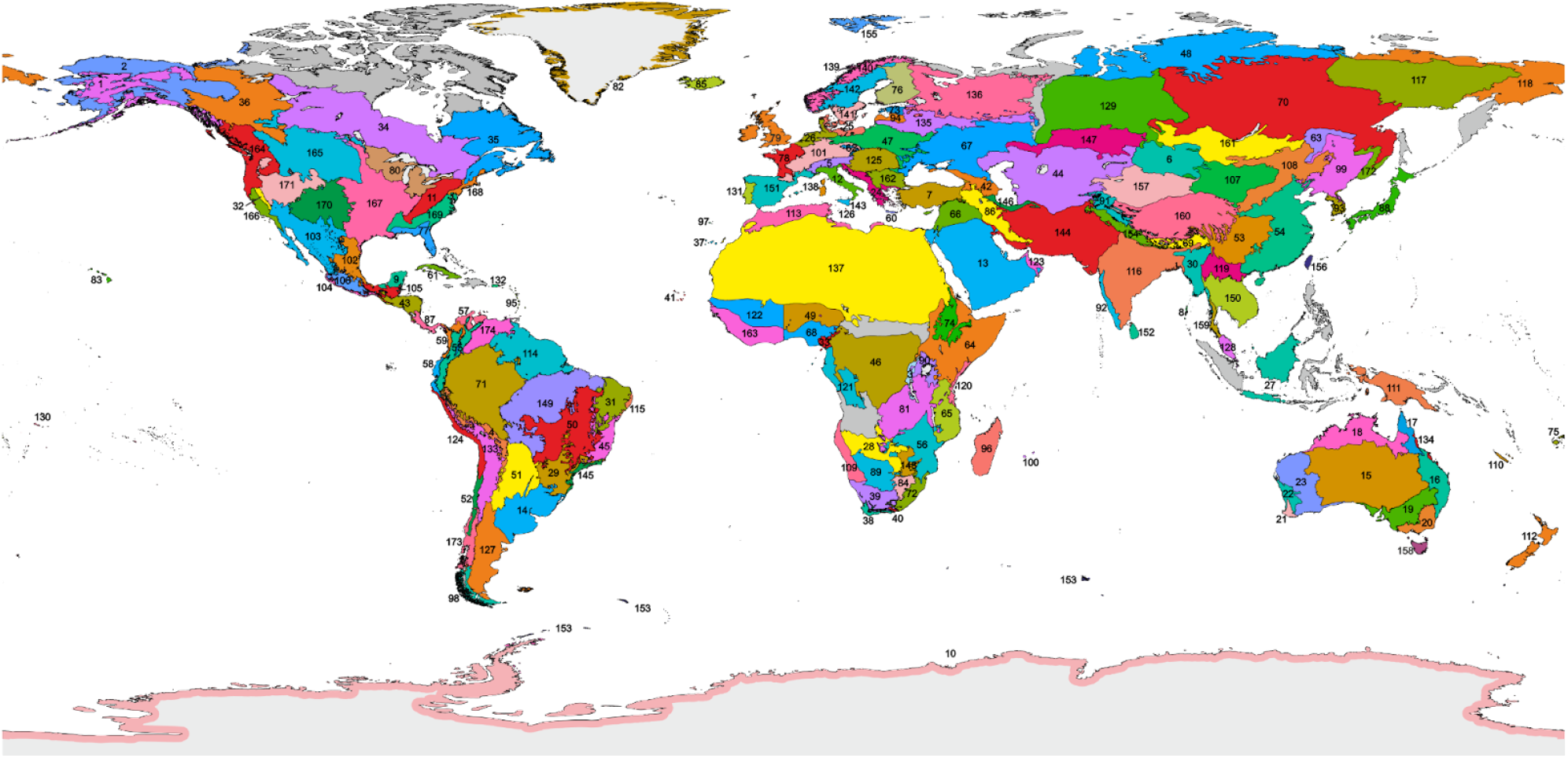
Distribution of 174 ecoregions used in endemicity analyses. Ecoregions excluded from the analyses due to the lack of data are indicated in gray. Their explanation is given in Table 1.

**Fig. 8.**
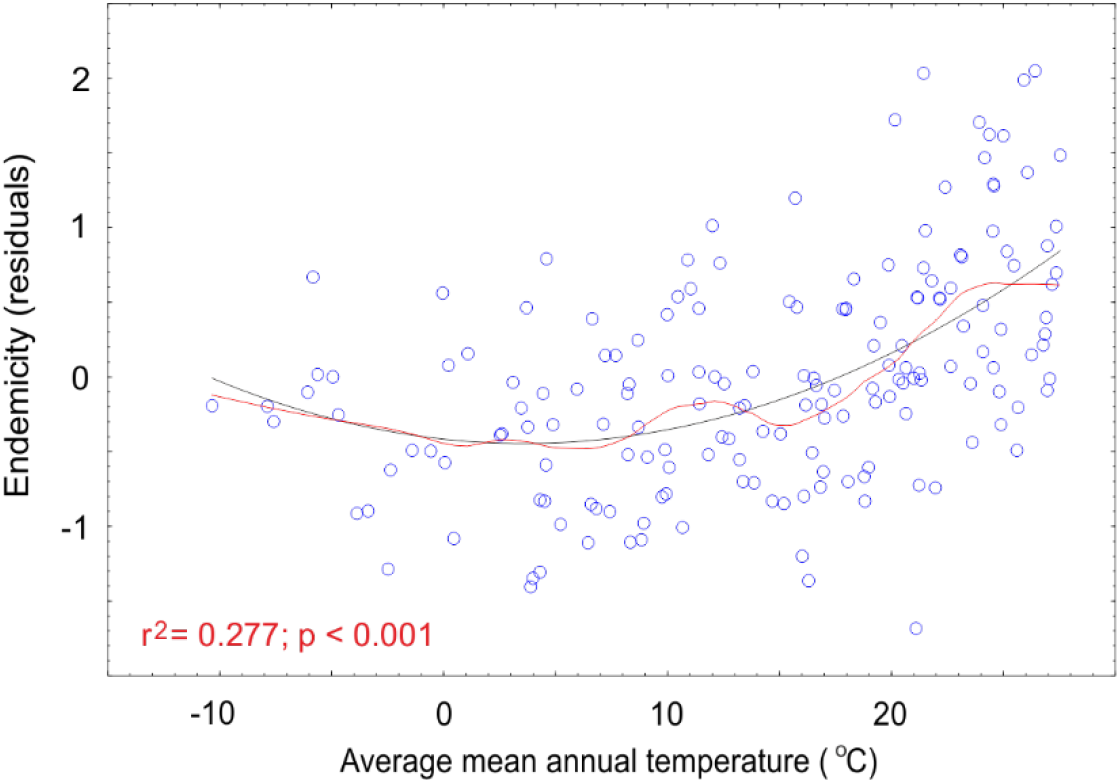
The effect of average mean annual precipitation on endemicity of fungi at the ecoregion scale. **B**lack line, best quadratic fit; red line, lowess curve.

#### Box 1. Calculation of endemicity indices and correlation among standardized indices for all fungi and among functional groups.

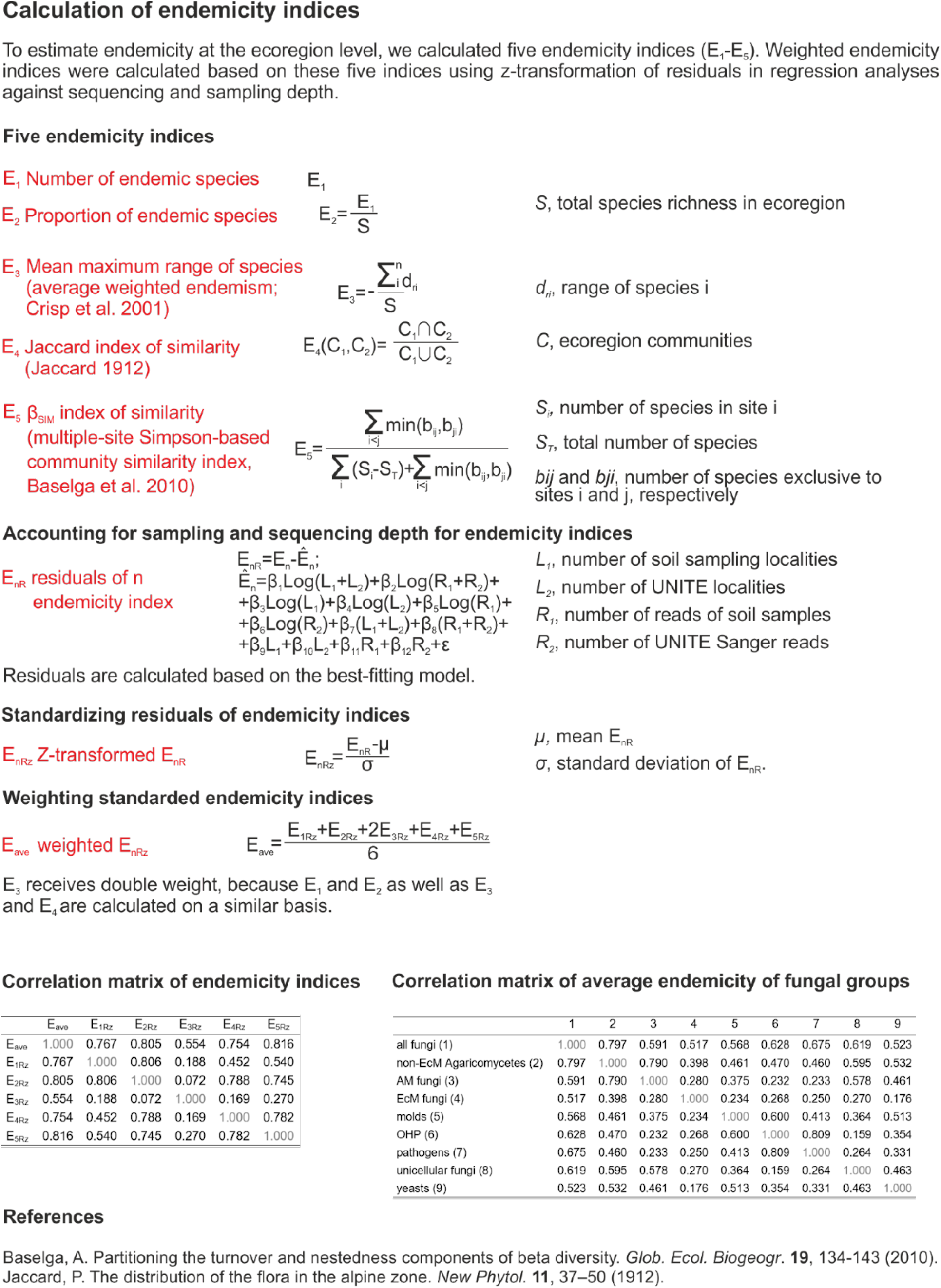

While endemicity patterns of non-EcM Agaricomycetes and AM fungi were similar to those shown for all fungi, different patterns were found for other functional groups. Endemicity of EcM fungi was related to high mean annual precipitation (MAP) (R^2^_adj_=0.147). Molds, pathogens and yeasts showed multiple endemicity hotspots. Molds (R^2^_adj_=0.199) and pathogens (R^2^_adj_=0.105) had relatively greater endemicity in strongly acidic or alkaline soils, indicating that extreme soil conditions may support unique soil biota, with limited effective dispersal across edaphically extreme habitats. Human footprint (see Methods) had a weak negative effect on endemicity of all fungi (R^2^_adj_=0.018), pathogens (R^2^_adj_=0.015), and OHPs (R^2^_adj_=0.056), suggesting that anthropogenic habitat loss or homogenization may affect endemic species (Finderup Nielsen et al. 2019). European ecoregions had the lowest endemicity for all fungi (R^2^_adj_=0.065), pathogens (R^2^_adj_=0.086) and unicellular fungi (R^2^_adj_=0.035) compared with those of other areas. Averaged current aerial bioclimatic variables better explained endemicity compared with the ranges of those variables or bioclimatic variables of soil and last glacial maximum (LGM). Climate change since the LGM had a weak positive effect on endemicity of molds (mean diurnal range and overall climate change: R^2^_adj_=0.073) and OHPs (isothermality and mean diurnal range: R^2^_adj_=0.056) but not other groups.

We found that patterns in fungal endemicity were relatively consistent among the five individual endemicity indices and that they resemble endemicity patterns of vascular plants and animals, which exhibit major hotspots in wet tropical habitats (Kier et al. 2009; Barlow et al. 2018). However, endemicity patterns in fungi were somewhat weaker, which may reflect the greater long-distance dispersal capacity of fungal spores relative to propagules of plants and animals (Golan & Pringle 2017). In terms of the greater macroorganism richness and endemicity found in the tropics, the literature abounds with hypotheses, including narrower niche breadth, more asymmetric interactions (i.e., greater specialization), climatic stability, and more rapid evolution due to environmental energy (Vazquez & Stevens 2004; Brown 2013) in the tropics than elsewhere. Negligible effects of the LGM suggest that climatic stability is not an important driver of fungal endemicity, a pattern that contrasts with those of plants and animals (Rosauer & Jetz 2015). The greater phylogenetic diversity of fungi noted for the tropics (e.g. Tedersoo et al. 2018) may in part reflect tropical origins for many lineages, as well as the radiation and rapid speciation of a limited number of EcM fungal genera into higher latitude areas (Kennedy et al. 2012; Sanchez-Ramirez et al. 2015). On a global scale, plant diversity does not appear to be causally related to fungal diversity (Tedersoo et al. 2014), but there is some evidence for stronger mutualistic plant-fungal interactions related to high rainfall (Põlme et al. 2018). Pathogenic interactions warrant further research in this respect, given their major importance as regulators of plant diversity (Chen et al. 2019). Tropical soil fungi have relatively greater dispersal limitations (Bahram et al. 2013) and narrower distribution ranges (Tedersoo et al. 2014), suggesting that high local diversity may contribute to greater regional-scale endemicity.

### Vulnerability of fungi to global change drivers

Communities with many species at their environmental niche limits may be particularly vulnerable to local extinctions (Watson et al. 2013; Smith et al. 2020b). Thus, we evaluated the relative vulnerability of soil fungal functional groups by estimating the percentage of species occurring at their upper niche limits to three major global change drivers – land use (land cover change), heat (maximum monthly temperature), and drought (lowest quarterly precipitation). We projected to the year 2070 relative to the 2015 baseline, using the average vulnerability index (Smith et al. 2020b), land use extrapolations of the LUH2 global dataset (Hurtt et al. 2020), and climatic extrapolations based on the CCS8.5 scenario (Karger et al. 2021). For all fungi taken together, predicted vulnerability to heat (best predictor: maximum monthly temperature; R^2^_adj_=0.583) and drought (precipitation seasonality; R^2^_adj_=0.456) were the greatest in the tropical and subtropical latitudes. Vulnerability to land use change (isothermality; R^2^_adj_=0.145) peaked in the tropics. The overall additive global change vulnerability was thus the highest in densely populated tropical and subtropical regions. Fungal functional groups had similar vulnerability patterns, which were mostly related to temperature. Among fungal groups, average vulnerability scores were highest for AM and EcM symbionts and unicellular fungi, but these scores differed only slightly across the global change drivers (**Fig. 9**). The actual vulnerability is probably underestimated for biotrophic pathogens and EcM fungi, because these groups associate with a limited number of plant species and are sometimes host-specific (Kennedy et al. 2015). Therefore, the loss of one of the few key symbiotic partners may greatly reduce the biotic niche of specialist fungi.

**Fig. 9.**
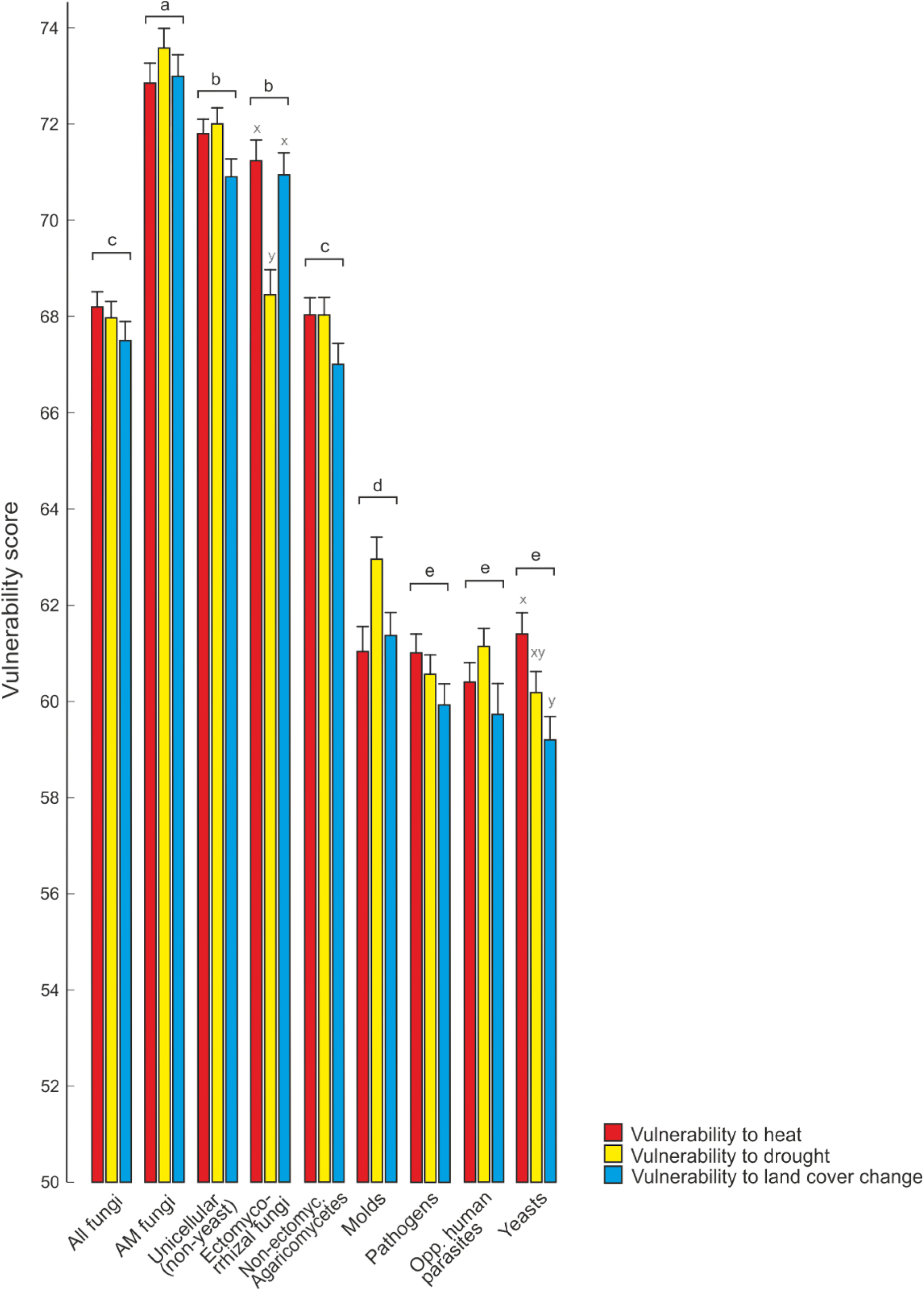
Vulnerability of fungi and functional groups to global change drivers. Different letters indicate statistically significant (*P*<0.001) differences among functional groups (a-e) and among global change drivers within functional groups (x-z).

Patterns of vulnerability in fungi are somewhat similar to those of terrestrial plants and animals, where vulnerability peaks in drylands prone to desertification (Warren et al. 2013), arctic/alpine areas (cold-adapted species), and regions with dense human populations (Watson et al. 2013). The relatively low vulnerability to heat in tundra-inhabiting fungi can be explained by their relatively high temperature optima (Maynard et al. 2019; but see Misiak et al. 2021), acclimation (Romero-Olivares et al. 2017), and poleward migration potential, despite relatively greater predicted warming in Arctic ecosystems. Above certain tolerance thresholds, soil organisms may be physiologically constrained by increasing soil temperature and evaporation, lower soil water potentials, and loss of oxygen due to greater respiration and faster decomposition, which result in hampered soil functioning and ecosystem multifunctionality (Delgado-Baquerizo et al. 2017). Open areas are predicted to increase due to climate change and human activities. This will further expose soil to solar radiation and result in the loss of fungal plant hosts. While here we calculated average vulnerabilities by adding up the effects of individual drivers, global change impacts tend to be synergistic (Rillig et al. 2019), so actual vulnerabilities may be much higher.

### Implications for conservation

Most fungi and soil organisms do not enjoy the protection and conservation measures that are afforded to more “charismatic” animals and plants (Ducarme et al. 2013). Nonetheless, fungi and other soil biota are pivotal to soil health, nutrient cycling, water storage, food security, and many other ecosystem services. Their biodiversity should hence be brought to the center stage of global sustainability thinking and conservation planning. For example, these organisms should be factored in when selecting protected areas otherwise based on plant and animal conservation (Guerra et al. 2021a). The fact that many EcM and plant pathogenic fungal species are associated with specific host plants indicates that on the local scale it is not only the narrowly-distributed species but also unique biotic associations that require focused conservation measures. From the fungal perspective, it is particularly important to protect plant species that act as hubs in modules of biotic interaction networks, because these hub species typically associate with multiple, distinct fungal partners (Põlme et al. 2018). In other cases, certain unique plant species or higher taxonomic groups should be prioritized. For example, in southern South America, the drought-sensitive tree family Nothofagaceae is the only group known to support EcM fungi that are endemic to this area (Godoy & Marin 2019)

Although the vulnerability to environmental change differed among fungal groups, their overall global patterns were similar. This suggests that broad habitat conservation measures may work for most fungal groups, including macroscopic non-EcM Agaricomycetes and EcM fungi as well as more cryptic pathogens and other groups. To accomplish this, fungi need to be incorporated into conservation frameworks (Gonçalves et al. 2021). Actions to fill existing information gaps at the local and global levels must also be taken, and global-scale surveys should take into account the soil biodiversity assessments, complementing the traditional collections-based assessment with metabarcoding of environmental DNA. This applies to national conservation evaluation programs and engagement in global policy-making initiatives, such as the System of Environmental Economic Accounting of the United Nations, World Biodiversity Forum, and Post-2020 Global Biodiversity Framework. Furthermore, promoting the red-listing of endangered fungal species at the national and global levels is critical (FAO 2020; IUCN 2021). Fungi need active and specific inclusion in national and global conservation policies and strategies, not just passive and implicit protection.

Our study provides evidence that soil fungi may be highly vulnerable to global change, which needs to be considered when planning how to preserve these key organisms in a changing world. As with plants and animals, fungi appear to be environmentally sensitive due to the strong impacts that land cover change, low moisture, and high temperatures have on taxonomic and functional composition (Brinkmann et al. 2019; Makiola et al. 2019; this study). The endemicity of fungi is highest in tropical forest biomes (Kier et al. 2009; this study), so conservation measures advocated for tropical plants and animals (Brooks et al. 2006; Barlow et al. 2018) are likely to conserve fungi. Tropical forests are under continued threat from deforestation and degradation driven by expanding agriculture, extractive industries, and infrastructural projects (Bebbington et al. 2018). Conservation of herbaceous wetlands, tropical rainforests and tropical woodlands is supported by our global fungal conservation priority map that accounts for endemicity, vulnerability, and γ-diversity (**Fig. 10**). Additionally, given the importance of soil pH for soil microbial diversity and composition, it is essential to prioritize areas with high pedodiversity or mixed landscapes including bogs, various forest types, and grasslands. As a crucial measure, desertification and loss of soil organic matter needs to be controlled by reducing the conversion of primary forest to crops and pasture (Smith et al. 2020a). This is important not only to prevent land degradation processes from impairing bacterial and fungal diversity, but also to sustain the capacity of drylands to provide essential functions and services, such as soil fertility, carbon storage, and food production for more than one billion people (Sivakumar 2007; Delgado-Baquerizo et al. 2018).

**Fig. 10.**
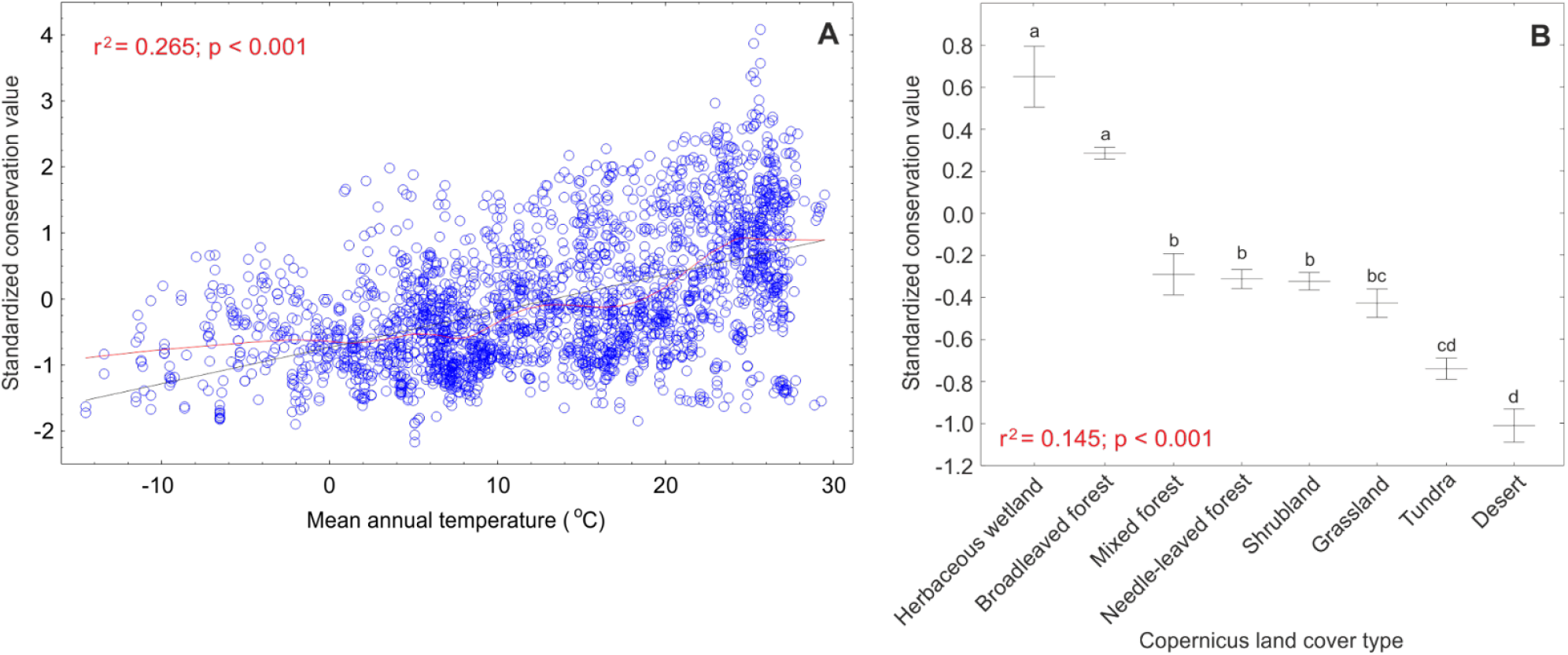
Relationships between conservation priority areas with mean annual temperature (A) and Copernicus land cover types. **(B)**. In A, black and red lines indicate best-fitting linear and lowess functions, respectively. In B, central lines and whiskers indicate mean and standard errors, respectively; letters above whiskers indicate statistically significant differences among land cover types.

## Conclusions

In conclusion, soil fungi show strong endemicity patterns, which differ by functional groups and are driven by both climatic and edaphic factors. Fungal groups also differ strongly in their relative vulnerability scores to global change, which peak in heavily populated tropical dryland areas. Unfortunately, these are the very areas most prone to further land degradation and desertification. Fungal endemicity and vulnerability patterns only partly mirror those of vascular plants and animals, which may be ascribed to their more efficient dispersal mechanisms. Global conservation efforts should include fungal biodiversity surveys alongside assessments of soil health, below-aboveground feedbacks, and areas of highest conservation priority, to secure the protection of habitats. Even more, they should include the monitoring of regional fungal communities over time, to pick relevant changes and to provide early warning signals of impending change. What we do not know, we cannot efficiently manage or protect.

## Methods

### Datasets

To study fungal endemicity and vulnerability to global change, we combined data from the Global Soil Mycobiome consortium (GSMc) open dataset (Tedersoo et al. 2021b) with materials from five other global soil biological surveys (**Fig. 1**) – BIODESERT (Maestre et al. 2022), MUSGONET (including the natural sites in Delgado-Baquerizo et al. 2021), CLIMIFUN (Bastida et al. 2021), GlobalAM (Davison et al. 2021), GlobalWetlands (Bahram et al. 2022) as well as Sanger sequence data from soil-inhabiting fungi obtained from the UNITE database (Nilsson et al. 2019) covering GenBank. We obtained the DNA from all five surveys and performed new DNA metabarcoding analyses following the protocols outlined for the GSMc dataset (Tedersoo et al. 2021b).

All datasets comprised information on geographical coordinates and soil pH. Based on geographical coordinates, we assigned the following climatic and land cover metadata to the samples: i) CHELSA v2.1 bioclimatic variables for the period 1981-2010 (Karger et al., 2020), ii) CHELSA-TraCE21k v1.0. for the LGM (Karger et al. 2021), and iii) CHELSA v2.1 climate extrapolations for the year 2070 following the RCP8.5 global warming scenario with SSP5 socioeconomic conditions and the GFDL-ESM4 global circulation model (Karger et al., 2020); iv) normalized difference vegetation index (NDVI; Filipponi et al. 2018); v) SoilGrids v.2 soil pH from 0-5 cm depth (Poggio et al., 2021); vi) land cover type using Copernicus classification v.3 (Buchhorn et al. 2020) for the year 2015; and vii) human footprint index based on the Land-Use Harmonization (LUH2; Hurtt et al., 2020) or the year 2015 and 2070 extrapolation. Based on original descriptions of vegetation (age, cover, relative abundance of species, fire history) or remote sensing data (Google Maps), samples were assigned to biomes (Olson et al. 2001) and land cover types. Based on Z-transformed differences in present and LGM bioclimatic variables, we calculated for each sample an averaged LGM climate change index. Further, for each sample we estimated the human footprint index as the cumulative sum of land-use state transitions, with the year 1960 used as a baseline.

### Bioinformatics

To infer fungal species and taxonomy, we used a long-read sequencing approach involving the ribosomal RNA 18S gene V9 subregion, ITS1 spacer, 5.8S gene, and ITS2 spacer to enhance taxonomic resolution and accuracy. We used degenerate, universal eukaryotic primers to cover as many divergent taxa within the fungi and micro-eukaryotes as possible (Tedersoo et al. 2021a). The amplicon samples were prepared in 82 PacBio SMRTbell sequencing libraries and sequenced on 48 PacBio Sequel 8M SMRT cells. The obtained reads were quality-filtered, demultiplexed to samples, trimmed to include only the full-length ITS region, and clustered to operational taxonomic units (conditionally termed as species) at 98% sequence similarity, which roughly corresponds to species-level divergence. Taxonomy was assigned based on information from the 10 best BLASTn matches against the UNITE 9.1 beta dataset (https://doi.org/10.15156/BIO/1444285). The resulting species-by-sample matrices were manually checked library-wise for external and cross-contamination and rates of index switching artifacts. We excluded several samples for which we suspected contamination, and removed rare occurrences of dominant species using the following thresholds: abundances = 1 for species with total abundance of >99 and abundances = 2 for species with total abundance of >999.

Based on FungalTraits 1.3 (Põlme et al. 2020), species belonging to the kingdom Fungi were assigned to functional groups based on ecological or physiological characters: i) AM fungi (including all Glomeromycota but excluding all Endogonomycetes, because there is not enough information to distinguish AM species from free-living species); ii) EcM fungi (excluding dubious lineages); iii) non-EcM Agaricomycetes (mostly saprotrophic fungi with macroscopic fruiting bodies; iv) molds (including Mortierellales, Mucorales, Umbelopsidales, and Aspergillaceae and Trichocomaceae of Eurotiales and Trichoderma of Hypocreales); v) putative pathogens (including plant, animal and fungal pathogens as primary or secondary lifestyles); vi) OHPs (excluding Mortierellales); vii) yeasts (excluding dimorphic yeasts); and viii) other unicellular (non-yeast) fungi (including chytrids, aphids, rozellids, and other early-diverging fungal lineages). Other groups such as lichen-forming fungi were not considered, owing to their relative infrequency in soil across samples and ecoregions. Among these groups, mostly non-EcM Agaricomycetes and EcM fungi comprise many red-listed species of conspicuous conch-shaped, resupinate, or stipitate fruiting bodies and are hence considered to be of relatively higher conservation interest (Cao et al. 2021; IUCN 2021).

### Fungal diversity

To assess patterns in global fungal α-diversity, we first calculated the residuals of logarithmically-transformed fungal richness and richness of major functional groups by performing linear regression against the logarithm of sequencing depth. We also compared other approaches such as residuals from untransformed richness against square-root-transformed and log-transformed sequencing depth (Tedersoo et al. 2014), exclusion of singletons, Shannon index of diversity, traditional rarefaction (depth, 500 reads), and SRS-rarefaction (Beule & Karlovsky 2020) to 500 (minimum) or 3894 (median) reads (**Fig. 2**). Because the approach including singletons and log-log transformation for selecting residuals resulted in best-supported models (**Fig. 2**), we chose this approach for further analyses. Because the sampling protocols differed in sampling area size, number of subsamples and DNA extraction protocols, we included only the GSMc dataset in analyses of fungal richness and composition. The GSMc dataset furthermore featured information about vegetation (age, proportion of dominant plant taxa and mycorrhiza types, fire history) soil properties (C, N, P, K, Ca, Mg concentration), and sampling date. Based on geographical coordinates and sampling dates, we calculated geographic and temporal eigenvectors using the adespatial package of R (Dray et al. 2018). Phylogenetic eigenvectors were calculated for woody plant species composition by mapping the taxa to a multigene vascular plant phylogram (Qian & Jin 2016).

Because of metadata availability and comparability, we exclusively used the GSMc dataset (3200 composite samples by 722,682 species) to estimate the best predictors of fungal α-diversity and composition and to update global fungal diversity distribution maps. We first calculated the residuals of logarithmically-transformed fungal richness and richness of major functional groups by performing linear regression against the logarithm of sequencing depth. Besides richness residuals and Shannon index of diversity, we calculated fungal phylogenetic diversity (PD), mean phylogenetic distance (MPD), and mean neighbor taxonomic distance (MNTD) indices based on a classification tree (Tedersoo et al. 2018) using the PhyloMeasures package of R (Tsirogiannis & Sandel 2016). For predictors, we included biome and continent (dummy variables), bioclimatic, edaphic, and vegetation-related variables, as well as eigenvectors of woody plant phylogenetic composition, spatial and temporal distance. Using random forest, we retrieved 20 candidate variables for GLM modeling. In GLM modeling, we included quadratic terms to account for non-linearity. To avoid an excessive number of predictors, only significant variables (P<0.001; R^2^>0.020) were kept in the final models. We also performed additional correlation analyses to illustrate the latitudinal gradient of α-diversity and γ-diversity. For generating fungal diversity maps, we performed additional analyses using bioclimatic variables, database pH_H20(0-5 cm)_, human footprint index and Copernicus land use categories and their interactions with continuous predictors as described for vulnerability maps. The GSMc dataset-based α-diversity analyses were validated by performing similar analyses using other datasets. Richness residuals were calculated separately for these data and all datasets were subjected to model selection for soil pH and bioclimatic variables, because information about other predictors was inconsistent. We also modeled various versions of soil pH including the original measurements of pH_KCl_ as well as pH_H20_ extrapolations for soil depths of 0-5 cm, 0-30 cm, and 15-30 cm. Since the pH_KCl_ fit significantly better into global models (**Fig. 2B**) and pH_H20_ extrapolations were missing for smaller islands and Antarctic habitats, we used pH_KCl_ in subsequent analyses.

For γ-diversity, logarithmically-transformed cumulative ecoregion (see below) species richness was subjected to model selection against the logarithmically-transformed number of samples and sequencing depth to calculate residual richness. Residual richness and endemicity indices of all fungi and functional groups were subjected to random forest machine learning analysis to pre-select ten most important variables for GLM.

### Endemicity

To infer endemicity patterns in fungi, samples (including data obtained from UNITE, covering International Nucleotide Sequence Databases entries) were assigned to ecological regions (Olson et al. 2001) based on their geographical coordinates, allowing a 10-km of buffer zone between a terrestrial ecological region and water (due to low resolution of the map layers in shore areas). Based on climatic and floristic similarities, the ecological regions were further aggregated into larger areas or split into smaller, geographically distinct units, which we refer to as ecoregions (**Fig. 7**). Each of 174 ecoregions comprised 1 to 45 soil samples, with surplus samples excluded randomly. Five indices of endemism – *viz*., the number of endemic species (weight = 16.7%), proportion of endemic species (weight = 16.7%), mean maximum geographical range of taxa (weight = 33.3%), Jaccard index (weight = 16.7%), and beta-sim index (weight = 16.7%) – were selected for calculating the averaged endemicity index based on the community matrix (Crisp et al. 2001; Villéger & Brosse 2012; **Box 1**). The former two and latter two indices reflect similar aspects of endemicity and were therefore downweighted. To account for differences in sampling intensity, we calculated residuals for the numbers of all species and endemic species by regressing these against the logarithmically-transformed number of samples and sequencing depth.

Of the indices used, only the number and proportion of endemic taxa were significantly positively correlated with species richness (all fungi: r=0.707 and r=0.212, respectively), whereas others had no significant correlation. Furthermore, species richness was not included among the best predictors of averaged endemicity, indicating that these metrics are independent. Endemicity indices were calculated using the betapart package v.1.5.4 (Baselga & Orme 2012) of R v.4.1.10 (R Core Team 2022). Endemicity indices of all fungi and functional groups were subjected to random forest machine learning analysis to pre-select the ten most important variables for GLM. We used the variance (as coefficient of variation) and averaged values of bioclimatic variables, area, latitude, longitude, altitude, and soil pH as well as continents (dummy variables) to explain endemicity. GLMs were fitted using second-order polynomial terms for continuous variables. Only significant variables (P<0.050; r^2^>0.020) were kept in the final models. Based on the predictions revealed by GLMs, endemicity maps were constructed using the sf v.1.0-5 (Pebesma 2018) package of R.

### Global change vulnerability

The vulnerability of soil fungal groups was estimated relative to three global change drivers – heat (maximum monthly temperature), drought (negative of inverse hyperbolic sine-transformed precipitation in the driest quarter), and land cover change – for the year 2070 (relative to 2015 baseline) using the community-mean percentile vulnerability index (V_2_; Smith et al. 2020b). This index is based on averaging percentiles of all species at a given global change driver value.

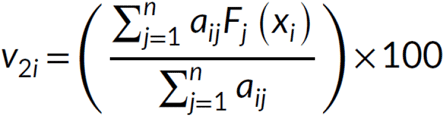

where a_ij_ is the presence (0 or 1) of species j in site i, F_j_(x_i_) is the percentile of species j given site parameter value x_i_, x_i_ is the parameter value of site i, and n is the total number of species observed.

Precipitation in the driest quarter was selected as a proxy for drought, because bioclimatic variables cover larger areas (including islands) and offer greater resolution compared with other measures of soil water content and indicators of drought. The vulnerability scores were calculated for each soil sample using vuln v.0.0.05 (Smith et al. 2020b) package of R. We also constructed the average vulnerability score by equally weighting all components. The vulnerability scores were unrelated to sequencing depth and sample size. We performed a similar random forest and GLM modeling exercise for determining the main predictors of vulnerability as described above, but allowed interaction terms between categorical and continuous predictors and used a more relaxed threshold for keeping variables in the model (P<0.001; R^2^>0.01) due to greater sample size. To construct vulnerability maps we used a regression-kriging approach (Hengl & MacMillan 2019). To predict vulnerability scores for each global driver and estimate their prediction uncertainty, thin plate splines (basis dimensionality = 3) were fitted using a generalized additive model (GAM) with mgcv v.1.8-38 (Wood 2011) package. To incorporate the spatial autocorrelation signal, we calculated residuals at the sampling sites and used inverse distance weighting (IDW) to interpolate residuals beyond the sampling sites. To obtain final vulnerability predictions, interpolated residuals were added to the results based on the predicted regression part. By using the relative vulnerability values, we also prepared the map of fungal vulnerability ascribed to each of the three components. Vulnerability maps were visualized using the raster v.3.5-9 (Hijmans 2021) package of R.

The maps for conservation priorities were calculated for all fungi using sampling points used in vulnerability analyses, except points corresponding to cropland and urban and village land cover. For each sampling point, the respective average endemicity, γ-diversity, and vulnerability scores were z-transformed, followed by adding a constant (5, to exclude negative values), multiplied (to downweigh areas with any low values), and used in a regression-kriging approach (**Table 3**) as described for vulnerability.

**Table 3.** The best models of conservation priority co-kriging maps for fungi. All P-values are <0.001.

|  | Df | Sum of squares | Mean squares | F-value | R<sup>2</sup><sub>adj</sub> | Trend |
| --- | --- | --- | --- | --- | --- | --- |
| air MAT<sup>1</sup> | 2 | 2133 | 1066 | 1426 | 0.266 | positive |
| MPWM | 1 | 992 | 992 | 663 | 0.134 | positive |
| climate isothermality | 1 | 169 | 169 | 113 | 0.023 | positive |
| MAP | 2 | 167 | 84 | 112 | 0.022 | positive |
| Land cover type<sup>2</sup> | 7 | 128 | 18 | 12 | 0.015 |  |
| Land cover type x MAP | 7 | 121 | 17 | 12 | 0.014 |  |
| error | 2477 | 3705 |  |  |  |  |

<sup>1</sup>Abbreviations: MAP, mean annual precipitation; MAT, mean annual temperature; MPWM, mean precipitation of wettest month;
<sup>2</sup>Copernicus land cover types: broadleaf forest, mixed forest, coniferous forest, shrubland, tundra, grassland, desert, wetland (note that cropland and urban and village biomes were excluded).

Based on methods comparison, we conclude that inclusion of singletons may increase richness model performance at relatively low sequencing depth in high-quality, third-generation sequencing datasets. Furthermore, residuals of log-transformed richness against log-transformed sequencing depth yields significantly better model estimates compared with most other standardization methods. Richness data from studies with different sampling designs must not be pooled in a common analysis unless the factor “study” or sampling attributes (e.g. number of subsamples, volume of samples) are accounted for. Use of original soil pH data strongly outperforms extrapolated data; therefore, soil pH extrapolations should not be used for testing relative effects of edaphic and climatic and other properties. This probably applies to other edaphic and vegetation-related values that greatly vary on a local scale.

## Acknowledgements

The bulk of the funding is derived from the Estonian Science Foundation (grants PRG632, PRG1170, PSG136, MOBTP198), Norway-Baltic financial mechanism (grant EMP442) and Novo Nordisk Fonden (NNF20OC0059948). LT, VM, AZ, MB, NAS, AA, UK, and KA designed the study; VM, LT, AZ, MB, NH-D, SA, and OP analyzed data; LT, MD-B, FTM, JP, MÖ, MM, MZ, ME, and ÜM contributed DNA extracts from global surveys; other authors contributed materials, data, and/or chemical analyses; LT wrote the first draft and all authors contributed to the writing of the article. The authors confirm that there are no competing interests. Data for this paper have been deposited to the PlutoF data repository (GSMc data: https://doi.org/10.15156/BIO/2263453. Representative sequences of identical reads per sample are available from the UNITE database. The scripts used for the bioinformatic analysis are available at GitHub: https://github.com/Mycology-Microbiology-Center/GSMc

## Notes

### Competing Interest Statement

The authors have declared no competing interest.

